# Scorpion α-toxin LqhαIT specifically interacts with a glycan at the pore domain of voltage-gated sodium channels

**DOI:** 10.1101/2024.01.26.577479

**Authors:** Swastik Phulera, Callum J. Dickson, Christopher J. Schwalen, Maryam Khoshouei, Samantha J. Cassell, Yishan Sun, Tara Condos, Jonathan Whicher, Wilhelm A. Weihofen

**Affiliations:** Discovery Sciences, Novartis Biomedical Research, 250 Massachusetts Avenue, Cambridge, MA 02139, United States; Computer-Aided Drug Discovery, Global Discovery Chemistry, Biomedical Research Novartis, 181 Massachusetts Avenue, Cambridge, MA 02139, United States; Discovery Sciences, Novartis Biomedical Research, Novartis Pharma AG, Basel, Switzerland; Neuroscience, Novartis Biomedical Research, 22 Windsor St, Cambridge, MA 02139, United States; Ignis Therapeutics (Suzhou) Limited, Suzhou, China; Johnson & Johnson Innovative Medicine, 301 Binney St, Cambridge, MA 02142, United States

## Abstract

Voltage-gated sodium (Nav) channels sense membrane potential and drive cellular electrical activity. Numerous protein toxins have been identified that modulate Nav gating, and structures of Nav channels in complex with these toxins helped elucidate the molecular mechanisms of voltage-dependent channel gating. The deathstalker scorpion α-toxin LqhαIT exerts a strong action potential prolonging effect on Nav channels. Biochemical studies show that LqhαIT features a functionally essential epitope at its C-terminus that is not shared with related scorpion α-toxins. To elucidate the mechanism of action of LqhαIT, we determined a 3.9 Å cryo-electron microscopy (cryo-EM) structure of LqhαIT in complex with the Nav channel from *Periplaneta americana* (NavPas). We found that LqhαIT binds to voltage sensor domain 4 and traps it in a “S4 down” conformation to stabilize the open state. To promote binding, the functionally essential C-terminal epitope of LqhαIT forms an extensive interface with the glycan scaffold linked to Asn330 of NavPas that augments a small protein-protein interface between NavPas and LqhαIT. A combination of molecular dynamics simulations, structural comparisons, and prior mutagenesis experiments demonstrate the functional importance of this toxin-glycan interaction. These findings help establish a structural basis for the specificity achieved by scorpion α-toxins and provide crucial insights for the development and optimization of new Nav channel modulators.

## Introduction

Nav channels are key drivers of cellular electrical activity. They are best known for initiating and propagating action potentials in the brain and other excitable tissues such as muscle and heart^1^. Nav channels are validated therapeutic targets for pain^2,3^, epilepsy, and various neurological and neuromuscular disorders^4,5^. Eukaryotic Nav channel α subunits are large, single-chain proteins with ∼2,000 amino acid residues, organized into four homologous domains (DI – DIV) and adopting a pseudo-tetrameric fold. Each domain consists of six transmembrane helices (S1-S6), with helices S1–4 constituting the voltage-sensing domain (VSD) and S5-6 forming the pore domain (PD) (Fig. 1A inset). The four PDs form a central channel with the VSDs arranged around it in a domain-swapped manner^1^. An intracellular linker connects the VSDs to the PDs between transmembrane segments S4 and S5 (S4-S5 linker). Within the VSDs, the S4 helix contains positively charged arginine residues (R1-5), serving as gating charges that cause the S4 helix to slide up and down in response to a changing membrane potential^1^. This S4 helix movement is regulated by a number of conserved residues on S2, including an aromatic residue known as the hydrophobic constriction site (HCS), which serves as an immobile reference point to track S4 helical movements across the membrane. At resting membrane potentials, the S4 helices move towards the intracellular side of the membrane, a conformation described as the resting state or “down” state^6^. Upon membrane depolarization, the S4 helices in VSD I–III move quickly towards the extracellular surface to adopt the activated conformation, repositioning the S4-S5 linker to open the Na+ pore. However, the movement of the S4 helix in VSD-IV towards the extracellular surface is about five times slower than that of VSD I-III. The eventual movement of the S4 helix of VSD-IV to this “up” conformation induces Nav channels to inactivate^1^. To date, all apo Nav channel structures, including NavPas from *Periplaneta americana* (American cockroach)^7,8^ and human Nav 1.1^9^,1.2^10^, 1.4^11^, 1.5^9,12,13^, 1.6^14^, 1.7^15^, and 1.8^16^ are broadly similar and adopt an inactivated state^17^.

**Figure 1.**
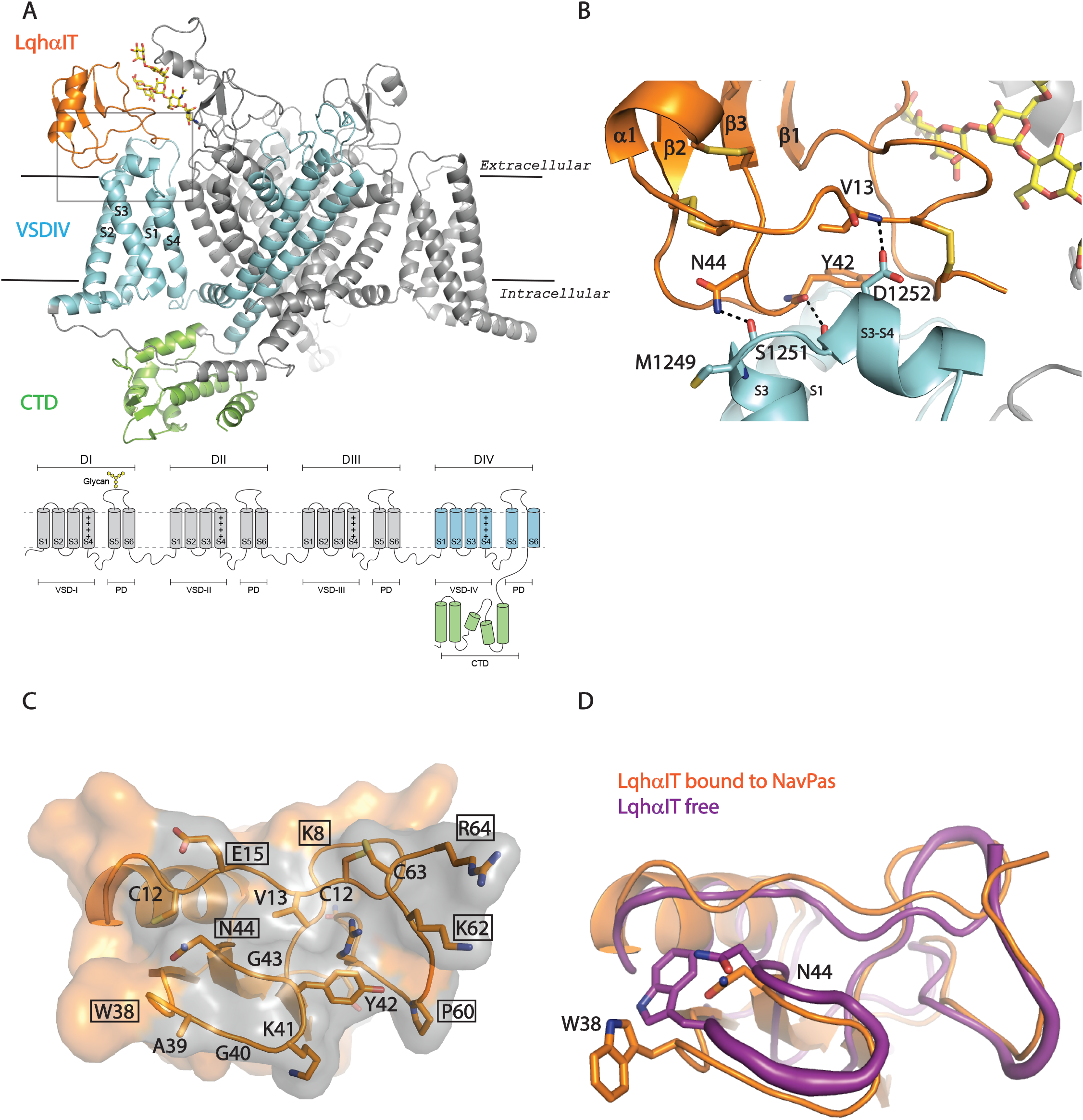
LqhαIT bound to the VSD-IV domain of NavPas. (A) VSD-III is hidden for clarity. LqhαIT is shown in orange, D-IV of NavPas in cyan, C-terminal domain (CTD) in green, and the remaining channel in grey. An Asn330-linked glycan chain (yellow) sandwiched between LqhαIT and NavPas is shown as sticks. Bottom Panel: Schematic of Nav channels. Asn330-linked glycan position on an extracellular loop of DI is shown. D-IV and the CTD are shown in cyan and green, respectively. (B) Polar interactions between LqhαIT and NavPas. Dashed lines indicate H-bonding interactions, side or main chain atoms of participating residues and disulfide bridges are shown as sticks. (C) The surface of LqhαIT is shown in orange and the interface with NavPas in grey. Residues lining the interface are shown as sticks. Residues analyzed by Karbat et al^25^ are boxed. (D) Structural/conformational changes in LqhαIT upon binding to the channel: Apo LqhαIT structure shown as a purple putty ribbon, channel bound LqhαIT structure in orange. Putty diameter reflects B factor distribution for the NMR structure of free LqhαIT (pdb code 1LQH). Side chains of Asn44 and Trp38 are shown as sticks.

Animal toxins exploit the excitatory role of Nav channels in different fashions. An important class of toxins that target Nav channels are scorpion α-toxins like LqhαIT and Aah2^18^. These toxins are typically >60 residues long and exhibit a knottin fold featuring an α-helix packed against a three-stranded β-sheet^19^. Four interlocking disulfide bonds further stabilize the fold. Previous structures revealed that scorpion α-toxins trap Nav channels in an activated Na^+^ ion conductive state, prolonging action potential duration by stabilizing the resting or “down” conformation of VSD-IV^13,20^. A-toxins share sequence similarity of ∼70% (Supplementary fig. 2D) and are classified based on their specificity for insect and mammalian Nav channels. For example, anti-mammalian toxin Aah2 from the scorpion *Androctonus australis Hector* and Lqh2 from the Middle Eastern scorpion, *Leiurus quinquestriatus hebraeus act* potently on mammalian Nav channels but exert negligible toxicity to insects^18^. In contrast, anti-insect α-toxins, such as LqhαIT from *Leiurus hebraeus* (Deathstalker scorpion) (60% sequence identity to Aah2), are very selective for insect Nav channels^18,19^.

**Figure 2.**
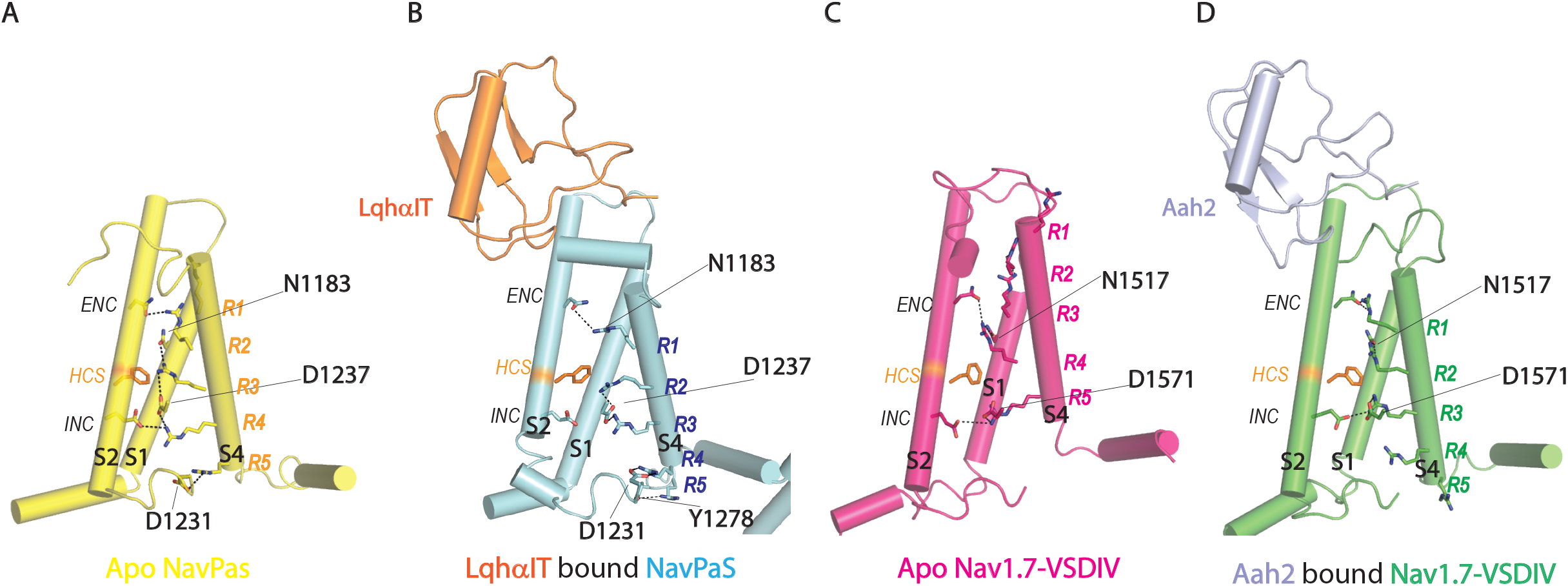
Activation and S4 helix movement in VSD-IV. (A) Side-view of VSD-IV of apo-NavPas. Side chains of extracellular negative charge cluster (ENC), intracellular negative charge cluster (INC), HCS, gating charges (R1-R5) as well as Asn1183 and Asp1237 which play a role in gating are shown as sticks, S3 not shown for clarity. (B) same as (A) showing LahαIT (orange) bound to NavPas VSD-IV. Also shown is Tyr1278 which interacts with R5 allowing the creation of an extra bulge at the end of S4, (C-D) same as (A) showing Nav1.7-VSD-IV apo and bound to Aah2 (blue), respectively.

Mutational studies show that Aah2 and LqhαIT share the same extracellular binding site on VSD-IV^21^, termed neurotoxin receptor site 3 and more recently as V4EM^22^. However, competition binding studies suggest that within receptor site 3, there are sub-regions differently targeted by toxin subtypes to achieve selectivity^23,24^. Specifically, two distinct epitopes on LqhαIT bind receptor site 3 and are essential for its function: a conserved “Core-domain” consisting of two loops harboring residues 17-18 and 38-44 and a variable “NC-domain” consisting of turn-forming residues 8–12 and C-terminal residues 56–64^25^. Swapping of NC-domain residues from LqhαIT onto the anti-mammalian α-toxin, Aah2, yielded a functional chimera with anti-insect activity^25^. However, the precise mechanism of how these residues determine the subtype preference of scorpion α-toxins remains unclear. Elucidation of the ability of various scorpion α-toxins to differentiate between insect and mammalian Navs, or among Nav subtypes in mammalian excitable tissues, is of potential value for the future design of selective drugs and insecticides.

To elucidate how the unique LqhαIT NC-domain epitope engages Nav channels and confers species selectivity, we set out to determine the structure of LqhαIT in complex with insect NavPas by cryo-EM. Our structure and subsequent molecular dynamics (MD) simulations demonstrate that LqhαIT interacts with VSD-IV and locks the S4 helix in a down conformation. We observed that functional residues within the NC-domain form specific interactions with a conserved extracellular glycan scaffold linked to Asn330 on the NavPas PD. These findings uncover how α-scorpion toxins achieve species selectivity between mammalian and insect Nav targets.

## Results

### Sample preparation and structure determination

To elucidate the mechanism of action of the anti-insect α-scorpion toxin LqhαIT on Nav Channels and the molecular basis for its species specificity, we determined the cryo-EM structure of NavPas bound to LqhαIT. NavPas^7^ and LqhαIT were expressed and purified separately. For complex preparation, NavPas was mixed with excess amounts of LqhαIT and subjected to size-exclusion chromatography (SEC) to isolate the complex (Supplementary fig. 1A). Non-uniform refinement in cryosparc yielded a final 3D reconstruction with an overall resolution of 3.9 Å (Supplementary fig. 1C). The local resolution gradually changes from ∼3.5 Å at the central pore region to about 5.5 Å at the most peripheral parts of the VSD domains (Supplementary fig. 1D). Compared to apo NavPas we observed additional density at the extracellular surface of VSD-IV, the expected binding site of LqhαIT (Supplementary fig. 1D). The additional density was resolved at 4 to 6 Å and exhibited sufficient structural features that allowed unambiguous placement of LqhαIT^26^ and define its interaction surface with NavPas. The refined LqhαIT-bound NavPas structure (Fig. 1A) is similar to the previously reported apo structure^7^ with an RMSD of 0.98 Å for 1037 superimposed residues. The permeation path for sodium ions along the pore is closed and calculated pore radii are comparable to the apo structure (Supplementary fig. 1E). There are however important local conformational differences observed in the two structures that are key to understanding LqhαIT’s mechanism of action.

### Conformational changes in VSD-IV

Nav channels embedded in detergent micelles do not experience a membrane potential, typically leading them to adopt an inactivated state. This state is characterized by the upward movement of the S4 helices in all four voltage sensors towards the extracellular side of the membrane to assume an “up” conformation^7^. In the inactivated state of apo NavPas two of the five arginine residues (R1 and R2) in the VSD-IV S4 helix are situated on the extracellular side of the HCS (Fig. 2A). The S4 adopts an α-helix around this R1-R2 segment and transitions to a 310-helix for the R3-5 segment located on the intracellular side of the HCS (Fig. 2A). Upon binding of LqhαIT, the S3-S4 loop, which lacks secondary structural elements in apo NavPas, becomes helical, and the S4 helix slides “down” one full helical turn to adopt a 310-helix throughout R1-R5 (Fig. 2B and Movie 1). The downward motion of S4 shifts R4 into a position to interact with Asp1231 in the S2-S3 loop, displacing R5. Concurrently, R5 moves into hydrogen bonding distance to Tyr1278 and the vicinity of phospholipid head groups (Fig. 2 A&B), and the S4-S5 linker, along with the intracellular tail of S5, undergoes a 3.5 Å displacement (Supplementary fig. 4). In this conformation of the S4, R1 remains on the extracellular side of the HCS, while R2 transitions to the intracellular side of the HCS (Fig. 2B). This suggests that a single gating charge transition across the HCS upon LqhαIT binding is sufficient to maintain NavPas in the open state. This contrasts with the structures of Nav1.7 and Nav1.5 in complex with Aah2^20^ and LqhIII^13^, respectively, where two gating charges traverse the HCS (Fig. 2, C and D). Furthermore, in both Nav1.5 and Nav1.7 apo structures there are three positively charged residues (R1-R3) extracellular to the HCS (Fig. 2, A and C), as opposed to NavPas, which has only two (R1-R2). Overall, our analyses demonstrate that the three V4EM targeting toxins trap respective S4 helices in comparable conformations, yet the channels’ inactivated states, characterized by the S4 helices in their most upward conformation, exhibit variations.

### VSD-IV-toxin interactions

Mutational studies demonstrated that the LqhαIT core domain epitope, comprised of residues in the β2-β3 loop and the β1-α loop, is essential for activity^25^. This epitope contacts the S1-S2 and S3-S4 linkers of VSD-IV in our structure (Fig. 1, A and B). We superimposed the channel-bound toxin with the NMR structure of the free toxin to identify any toxin regions that undergo conformational changes upon binding to NavPas (Fig. 1D). Most structural differences are within ∼1.5 Å RMSD except for residues 38-44 in the β2-β3 loop and residues 6-18 in the β1-α loop and the C-terminus. Within the β2-β3 loop, re-positioning of the Trp38 side chain is necessary to facilitate the H-bonding interactions between Asn44 and Tyr42, and Met1249 and Ser1251 of the NavPas S3-S4, respectively (Fig. 1 B). Mutating Trp38 and Asn44 resulted in a ∼9- and 30-fold loss in activity, respectively, demonstrating the importance of these core domain residues for toxin-channel interaction^25,27^. In the β1-α loop of LqhαIT the Val13 backbone forms a hydrogen bond with Asp1252 in the S3-S4 region of the channel (Fig. 1B). The C-terminus fills the aqueous cavity between VSD-IV and the S5 helix of the D1 pore domain, wherein the Arg64 side chain is inserted into a negatively charged cleft (Fig. 4B).

To validate the interactions in our model and identify additional interactions, we performed molecular dynamics (MD) simulations on the toxin-channel complex. For the simulations, the toxin-channel complex was embedded into a membrane bilayer consisting of 686 lipid molecules and equilibrated for 100 ns. From this starting point, we performed 10 replicate 200 ns production simulations (totaling 2 μs of sampling). The complex remained stable as reflected in Cα-RMSD fluctuations below 5 Å. We recorded the occupancies of all H-bonds (Fig. 3A) and identified eight hydrogen bonds (Fig. 3A) between the channel and toxin, persisting for at least 20% of the simulation duration. Importantly, all hydrogen bonds observed in our static EM structural model remained intact for at least 25% of the time. Furthermore, a hydrogen bond between Gly40 at the β2-β3 loop and the backbone of Leu1248 persists for over 80% of the time (Fig. 3A). We observed four salt bridges formed for at least 20% of the simulation time between toxin and channel (Fig. 3B). These involve Lys41 contacting Glu1200 and Asp1203 in the S1 helix, as well as Arg58 and Lys62 engaging with Asp1252 and Asp1255 in S3-S4 region, respectively (Fig. 3C). The salt bridge between Arg58 to Asp1252 is positioned at the center of the interface, shielded from solvent and other potential polar interaction partners. Notably, both these salt bridges appear critical, as conservative mutation of Arg58 to Lys or Lys62 to Arg reduces binding affinity by 77 and 11-fold, respectively (Fig. 4D)^25^.

**Figure 3.**
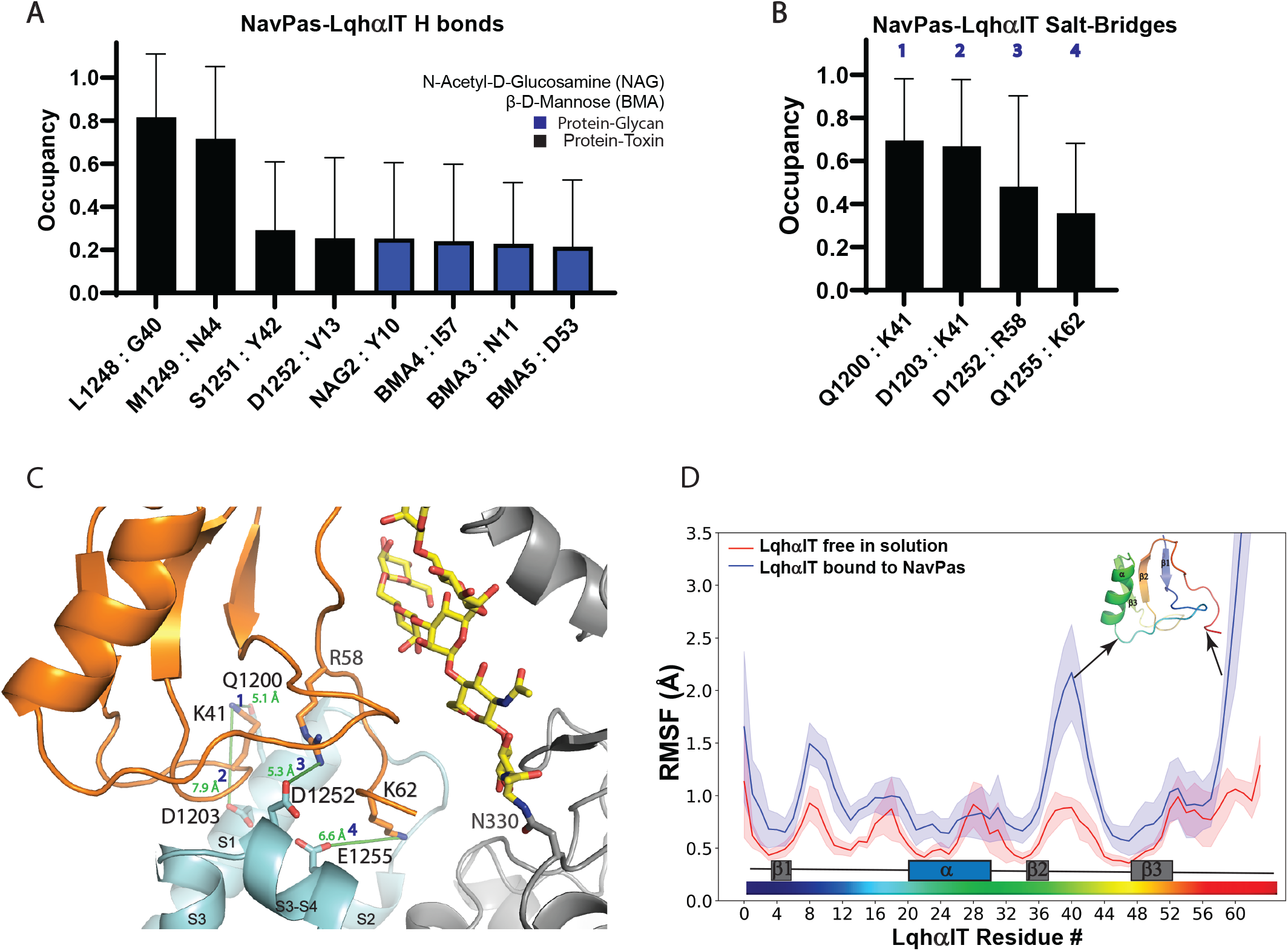
Channel-toxin interactions observed during MD simulations. (A) H-Bonds and (B) salt bridges during MD simulations. Data are expressed as the occupancy (% of frames) in which the interactions were present. (C) Frequently observed salt bridges (green lines) in (B) mapped to the cryoEM structure (NavPas VSD-IV is cyan, NavPAS PD is grey, Asn330 glycan is yellow, and LahαIT is orange). Blue numbers indicate the interacting pair in (B), distances in Å between residues in green. (D) Change in LqhαIT dynamics upon binding to NavPas. Cα RMSF values for free and channel-bound toxin during MD simulations. α helix and β sheets are indicated at the bottom. A significant reduction in RMSF values is observed for the β2-β3 loop and C-terminus.

**Figure 4.**
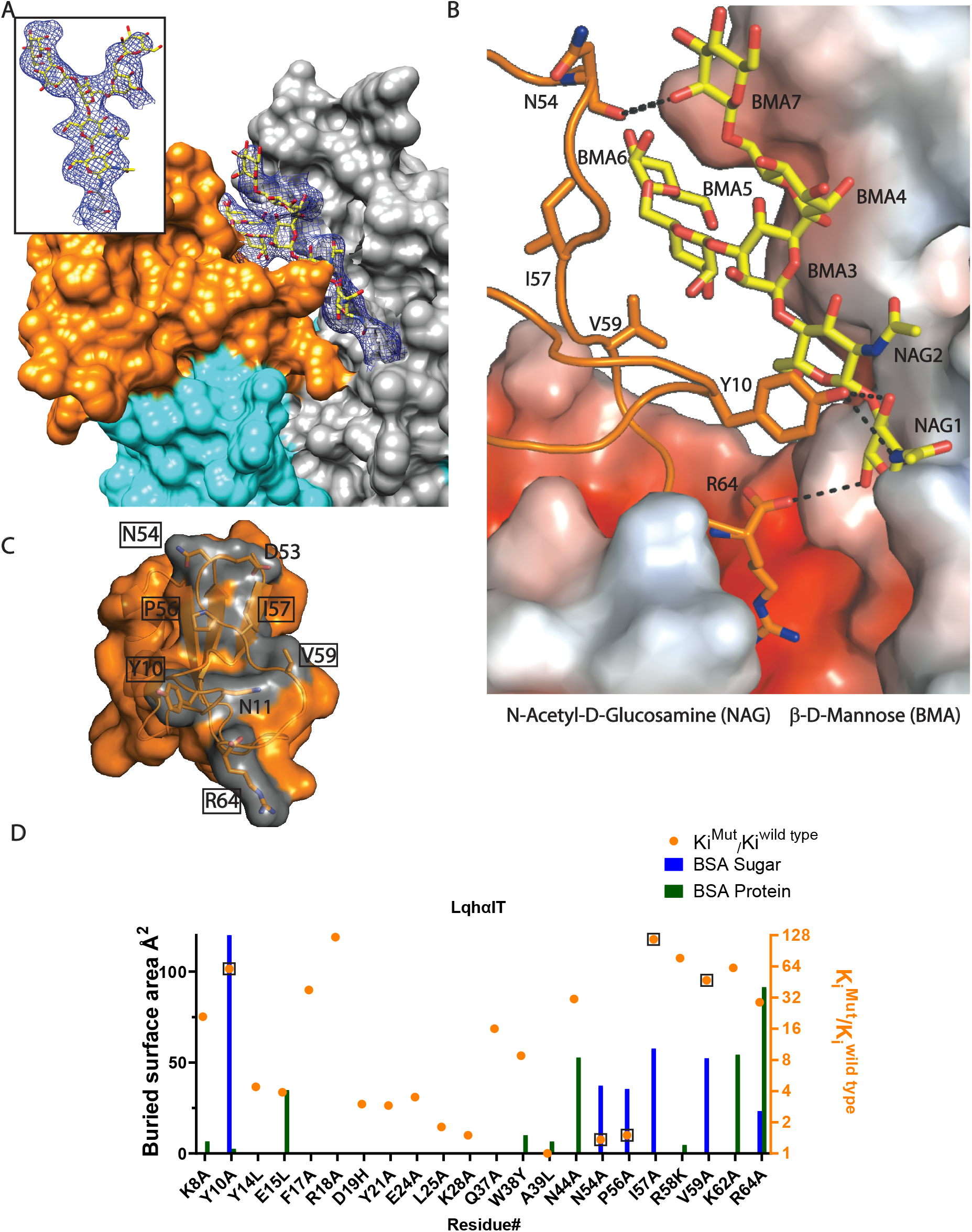
Toxin-glycan interactions. (A) Asn330-linked glycan chain sandwiched between LqhαIT and PD. Glycan residues (yellow) are shown in sticks surrounded by the meshed cryoEM map. Surface representation of toxin (orange) and channel (VSD-IV: cyan, rest of the channel: grey). (B) Toxin-glycan H-bonds and electrostatic interactions: Electrostatic charge on the channel surface is shown as calculated by APBS^62^ in pymol^63^, colored from a red to blue spectrum (−5 to +5 kT/e). Dashed lines indicate H-bonds between the toxin and glycan residues, additional residues that are in van-der-Waals distance to the glycan are shown as well. Arg64 from the toxin C-terminus is buried in the negatively charged pocket between VSD-IV and the PD domain. (C) LqhαIT surface patch that interacts with the glycan is shown in orange. Surface residues buried by the glycan in the toxin-channel complex are shown as sticks. Residues contributing to toxin function^25^ are boxed. (D) LqhαIT mutagenesis data from Karbat et al^25^ plotted against buried surface area changes upon complex formation calculated with PISA^64^. X axis indicates the amino acid mutation and the Y-axis on the right shows the corresponding change in binding affinity as well as the ratio of Ki values of mutant over WT (orange dots). Residues at the interface have their Ki values boxed. Blue bars represent the area buried by the glycan moiety while dark green bars represent the area buried by NavPas residues.

We also employed MD to investigate the dynamics of the free toxin and draw comparisons with the NavPas-bound state. Our MD simulations for the free toxin align with its dynamic features observed by NMR^26^, indicating heightened mobility in the β2-β3 loop and the unstructured C-terminus (Fig. 3D). Upon binding to NavPas, LqhαIT exhibits a reduction in backbone mobility, averaging 0.33 Å RMSF per residue. As anticipated, the β2-β3 loop assumes greater rigidity due to extensive interactions with the VSD-IV S3-S4 region of NavPas (Fig. 3, A and B).

Overall, the binding modes of LqhαIT and the related Aah2 are very similar, with both toxins adopting comparable poses when interacting with their respective Nav channel binding partner (Supplementary fig. 2A). Moreover, the shape complementarity of the contacting surfaces in each complex is akin: we calculated 0.421 for LqhαIT-NavPas complex vs 0.475 for the Aah2-Nav1.7 chimera complex using the shape correlation statistic (Sc)^28^. However, a notable contrast exists in the protein-protein interface sizes. The Aah2 and Nav1.7-VSD-IV interface measures 739 Å^2^, whereas it is only 495 Å^2^ between LqhαIT and NavPas. Despite this substantial difference, Aah2 and LqhαIT bind with comparable affinities to mammalian and insect ion channels, respectively, suggesting the potential contribution of an additional functional surface to LqhαIT’s affinity.

### Toxin-Glycan interactions are essential for insecticidal activity of α-scorpion toxins

Regarding the functionally essential NC-domain of LqhαIT, we observed no interactions with NavPas protein residues in both the EM structural model and subsequent MD simulations, except for the Lys62:Gln1255 salt bridge (Fig. 3C). Instead, the poorly conserved NC-domain epitope of LqhαIT establishes a substantial contiguous interface of 364 Å^2^ with the glycan scaffold linked to the conserved Asn330 in the D1 pore domain of NavPas (Fig. 4 A, B and Supplementary fig. 2D). This glycan chain, wedged between LqhαIT and the PD turret, is clearly defined in the cryoEM map (Fig. 4A), allowing us to confidently model seven monosaccharides. Seven out of the eight toxin residues lining the NC domain epitope exclusively engage with the glycan (Fig. 4C).

In the mutational study by Karbat et al^25^, five of these residues were analyzed, revealing that substituting Tyr10, Ile57 and Val59 resulted in a minimum 30-fold reduction in toxin activity. Tyr10 forms hydrogen bonds with the first two N-acetylglucosamines at the base of the glycan chain (Fig. 4B), while Ile57 and Val59 engage in van der Waals interactions, primarily with the mannose in the 1-3 arm (Fig. 4B). These findings indicate that LqhαIT interactions with the Asn330 glycan are highly specific, playing a crucial role in toxin affinity and specificity toward a glycan present at a conserved Asn330 position and a turret conformation that effectively restricts the mobility of the glycan through potentially specific interactions.

Glycan-toxin interactions were evident in the structure of Aah2 bound to a chimeric NavPas containing the VSD-IV from Nav1.7. Notably, the Aah2-Glycan interface (∼186 Å^2^) was approximately half the size of the LqhαIT-glycan interface (364 Å^2^), featuring a lone hydrogen bond between Asn9 and the second GlcNac unit (Supplementary fig. 2B). Intriguingly, despite this interaction, mutations at Asn9 in the related anti-mammalian toxin Lqh2 had no discernible impact on the toxin’s activity^27^. Moreover, mutations within the five-residue turn of Lqh2 (Asp8, Asp9, Val10) did not affect its function (Supplementary fig. 2E)^27^, whereas analogous mutations in LqhαIT (Lys8 and Tyr10) drastically disrupted insecticidal activity (Fig. 4D)^25^. The Asn330 glycosylation site is present in all mammalian Navs, except for Nav1.7 and Nav1.9. However, among the toxin/mammalian Nav complex structures available to date, none have revealed a toxin-glycan interaction (Supplementary fig. 3C). These observations strongly imply that the toxin-glycan interactions play a pivotal role in conferring selectivity of LqhαIT for insect over mammalian channels^25,27^.

## Discussion

The NavPas-LqhαIT complex structure presented here is the first documentation of an insect-specific scorpion α-toxin bound to an insect channel. This structure provides important insights into channel gating, the mechanism of channel modulation orchestrated by the toxin, and specificity exhibited by the toxin for insect over mammalian voltage-gated sodium channels. Notably, upon binding of LqhαIT to NavPas, the VSD-IV S4 helix shifts one helical turn towards the intracellular side in comparison to the Apo-NavPas structure. In the process a single gating charge transition across the HCS, signifying a Cα distance shift of ∼6.6 Å between two equivalent gating charge residues and roughly one 310-helical turn. This stands in contrast to the structures of Aah2 bound Nav1.7-VSD-IV and Lqh-III bound to Nav1.5, where the S4 was translated inwards by ∼14 Å and two of the gating charges transition across the HCS. This increased movement stems from a different inactive state conformation in mammalian Navs, placing the S4 helix one turn further towards the extracellular side.

Biochemical studies have suggested that anti-mammalian and anti-insect α-scorpion toxins might occupy distinct binding sites within receptor site 3 of Nav channels^29^. However, the driving force behind the high specificity of these toxins towards their respective Nav channel subtypes remained elusive. We combined previous mutagenesis data, cryo-EM structure determination and molecular dynamics, using the potent anti-insect scorpion α-toxin, LqhαIT, bound to NavPas to unravel the driving forces behind this specificity. We found that residues from the NC-domain form extensive contacts with the glycan chain attached to Asn330, thereby complementing the comparatively small interface between the core domain and VSD-IV (Supplementary fig. 2D). The unexpected close interaction with a protein glycan is noteworthy, as glycan chains rarely form specific, non-ionic interactions with protein binding partners. Previous biochemical and mutagenesis studies underscore the important role of the toxin/glycan interaction for the activity both anti-insect α-scorpion toxins and related sea anemone toxins ^25,27,30–32^.

The involvement of glycans in our structure may offer an explanation for the voltage independence of anti-insect α-scorpion toxins^33^. This is attributed to the extracellular surface of VSD undergoing conformational changes in response to voltage, while the PD glycan region likely remains stable even during voltage shifts. Additionally, the utilization of distinct binding principles by these toxins, each with its specificities (anti-insect vs anti-mammalian toxins), contributes to understanding the proposed shared macrosite concept^29^.

The challenges in advancing Nav channel modulators in clinical settings revolve around the difficulty in generating selective therapeutic compounds^5^. Conversely, natural product toxins, particularly peptides, can demonstrate both potency and selectivity, albeit presenting developmental challenges associated with the peptide modality^4^. LqhαIT’s binding serves as an example of this paradigm, showcasing both high potency and selectivity for insect over mammalian Navs^25^. Our cryoEM structure indicates that this results from a combination of protein-protein interactions and extended contacts to a posttranslational glycan modification absent in Nav1.7 and 1.9. This glycan-toxin interaction seemingly compensates for the smaller protein-protein interaction, which exhibits 33% less buried surface area when compared with AaH2. The ability to interact with the glycan present on insect Nav channels, as depicted in our structure, introduces another mechanism contributing to target selectivity.

## Methods

### Trx-LqhαIT expression and affinity purification

Oligonucleotides and E. coli codon-optimized genes were purchased from Integrated DNA Technologies Inc. (Coralville, IA). Cloning was performed using Gibson assembly with NEBuilder HiFi Assembly cloning kit according to the manufacturer’s specifications. Constructs were verified by Sanger sequencing at Genewiz (Cambridge, MA). *E. coli* SHuffle T7 Express (New England Biolabs, Ipswich, MA) cells were transformed with 100 ng pET32-Trx-LqhαIT and, cells were grown for 24 h on Lysogeny Broth (LB) agar plates containing 100 μg/mL carbenicillin at 30 °C. Single colonies were used to inoculate 10 mL of LB containing carbenicillin and grown at 30 °C for 20 h. This culture was used to inoculate 1 L of LB containing carbenicillin and grown to an optical density OD600 = 0.6. Protein expression was induced with the addition of 400 μM isopropyl β-D-1-thiogalactopyranoside (IPTG) for 16 h at 16 °C. Cells were harvested by centrifugation at 4700 rpm for 20 min, washed with phosphate-buffered saline (PBS), and centrifuged at 4700 rpm for 20 min. Cell pellets were stored at -80 °C until purification.

Cell pellets were resuspended in 30 mL lysis buffer (50 mM Tris-HCl pH 8.0, 300 mM NaCl, 15 mM imidazole, 0.1% Triton X-100, 2.5% glycerol (v/v)) containing 4 mg/mL lysozyme and 100 μL Protease Inhibitor Cocktail Set III (EDTA-free, MilliporeSigma). Cells were homogenized by sonication (3 × 30 s on ice with 10 min equilibration periods at 4 °C with rocking) using a Q500 sonicator (500 W) at 60% power (Qsonica, Newton, CT). Soluble protein was obtained by centrifugation at 10,000 rpm for 60 min at 4 °C. The supernatant was then loaded onto 3 mL of pre-equilibrated Ni-NTA resin (Thermo Fisher Scientific). The column was washed with 30 mL of lysis buffer followed by 90 mL of wash buffer (50 mM Tris-HCl pH 8.0, 500 mM NaCl, 20 mM imidazole, 2.5% glycerol (v/v)). His6-tagged Trx-LqhαIT was eluted with 30 mL elution buffer (50 mM Tris-HCl pH 8.0, 500 mM NaCl, 250 mM imidazole, 2.5% glycerol (v/v)) and concentrated/buffer exchanged with storage buffer (50 mM HEPES pH 7.3, 300 mM NaCl, 2.5% glycerol (v/v)) using a 10 kDa MWCO Amicon Ultra centrifugal filter (Millipore). Purified Trx-LqhαIT fusion protein concentration was determined by 280 nm absorbance on a Nanodrop 2000 spectrophotometer (Thermo) and stored at -80 °C.

### Trx-LqhαIT tag cleavage and HPLC purification

TEV protease was added to Trx-LqhαITs in 20 mol % in reaction buffer (50 mM Tris-HCl pH 7.5, 125 mM NaCl, 20 mM MgCl2 7·H_2_O) and reacted at 22 °C for 20 hr. TEV and Trx were removed from reaction mixtures by flowing over 2 mL of equilibrated Ni-NTA resin and washed with 5 mL of Ni-NTA buffer (50 mM Tris-HCl pH 8.0, 150 mM NaCl). Samples were desalted using a C18 HyperSep SPE cartridge (Thermo Fisher Scientific, 1 g bed weight), which was equilibrated with 10 mL MeCN + 0.1% formic acid (FA), 10 mL 50% MeCN aq. + 0.1% FA, and 20 mL 5% MeCN aq. + 0.1% FA. Loaded samples were washed with 10 mL 5% MeCN + 0.1% FA and eluted with 10 mL 80% MeCN aq. + 0.1% FA and dried under reduced pressure.

LqhαIT was purified using a Waters HPLC equipped with a Waters 2545 binary gradient module pumps, a Waters 2767 sample manager/fraction collector, Waters XSelect CSH Prep C18 preparatory column (19x150 mm), and a Waters Acquity QDa electrospray ionization (ESI) single-quadrupole mass analyzer. Samples were dissolved in 2 mL of 50% MeCN aq. + 0.1% FA and injected with a gradient mobile phase of water/MeCN + 0.1% trifluoroacetic acid (TFA, v/v) from 5-15% MeCN over 10 minutes at a flow rate of 30 mL/min. Elution was monitored by absorbance at 280 nm and by mass spectrometry. Fractions containing the LqhαIT were flash frozen in isopropanol/dry ice bath and lyophilized to dryness.

### Mass spectrometric characterization of Trx-LqhαIT

Trx-LqhαITs (100 μg) were TEV cleaved in a final volume of 100 μL and 10 μL were injected into a Waters Acquity UPLC system equipped with an ACE Excel 2 C18-Amide column (50 × 2.1 mm id, 2 μm particle size) and Sciex TripleTOF 6600 ESI quadrupole time-of-flight (QTOF) mass analyzer. TEV-cleavage reactions or purified Trx-LqhαITs were diluted to 0.1 mg/mL and 10 μL were injected with a gradient mobile phase of water/MeCN + 0.1% FA over 6 minutes from 0 – 30% MeCN at a flow rate of 0.6 mL/min.

### NavPas Protein expression and purification

For mammalian expression using HEK cells, we cloned a codon-optimized NavPas construct (based on the expressed protein sequence in^7^) in between the HindIII and XbaI sites of the vector BacMam pCMV Dest (Thermo Fisher Scientific). The. Bacmid was purified using the manufacturer’s instructions followed by transfection in Sf9 cells grown in suspension using ESF921 media (Expression Systems, CA) with X-tremeGENE HP transfection reagent (Roche). To make P2 virus 1 L of Sf9 cells in ESF921 media were infected with 1 mL of P1 virus. The culture was centrifuged and the supernatant containing P2 virus supplemented with 1% FBS before storage in the dark at 4 °C. The virus titer was determined by gp64-PE mouse anti-baculovirus antibody (Expression Systems, CA) using a Guava benchtop Flow Cytometer (Millipore, Sigma).

Protein was expressed using Expi293F cells grown in Expi293 Expression medium (Thermo Fisher Scientific). 2 L Cells were grown in 5 L Optimum growth flasks (Thompson, CA) at 110 RPM (orbital diameter of 50 mm) at 37 °C and 8% CO_2_. When cells reached a density between 2 and 3 × 10^6^ cells / mL, they were infected with P2 virus at an MOI of ∼1-1.2. 24 hrs. after infection at 37 °C, sodium butyrate was added to a final concentration of 10 mM, and the flasks shifted to 30 °C for another 48 hrs incubation. Cells were harvested by centrifugation and the pellets were stored at -80 °C.

Cells from 2 L culture were suspended in TBS, solubilized by adding 1% LMNG, 0.1% CHS, 1X cOmplete, EDTA-free protease inhibitor cocktail (Roche) and stirred for 1 hr. at 4 °C. Insoluble debris was pelleted by ultracentrifugation at 125,000 G for 45 min, and the supernatant containing the solubilized protein was collected for affinity purification by batch-binding to 10 mL of M2-agarose FLAG resin (Sigma) for 1 hour at 4 °C. Unbound proteins were washed with 9 column volumes (CV) of purification buffer (25 mM Tris pH 7.5, 50 mM NaCl, 0.1% (wt/v) digitonin. The protein was eluted with 5 CV of purification buffer supplemented with 150 μg/mL 3X-FLAG peptide (Apexbio). Fractions containing NavPas were pooled and concentrated. A 3-fold molar excess of LqhαIT and 1mM Veratridine (VTD) was added, and the mixture incubated on ice for 30 mins followed by size exclusion chromatography (SEC) on a Superose 6 increase column (Cytiva, MA) equilibrated in purification buffer. The peak fraction was concentrated and spiked with VTD to a final concentration of 1mM and a molar equivalent of LqhαIT to fully saturate the complex.

### Cryo-EM grid preparation and data collection

Electron microscopy grids (Quantifoil Au, 200-mesh copper R1.2/1.3) were glow-discharged for 90 s using PELCO easiGlow. 3.5 μl of sample was applied on a grid in the Vitrobot Mark IV (Thermo Fisher Scientific) chamber set to 100% humidity at 4 °C. The sample was blotted for 4 secs with a blot force of 25 and plunged into liquid ethane.

Datasets were collected on a Thermo Fisher Scientific Titan Krios microscope operated at 300 kV (FEI) and equipped with Falcon-IV direct electron camera (Thermo Fisher Scientific). Movies were taken in nanoprobe mode, with 50-μm C2 aperture and pixel size of 0.66 Å. Each movie comprises 50 subframes with a total dose of 50 e− per Å^2^. Exposure time was 3.08 s with a dose rate of 7.60 e^−1^ pixel^−1^ s^−1^ on the detector. Data acquisition was done using EPU software (Thermo Fisher Scientific) at −500 nm to -1.5 μm defocus (200 nm steps). Fringe Free Illumination (FFI) allowed four acquisitions per hole.

### Image processing

A total of 10,185 movies were collected and subjected to motion correction by MotionCorr2^34^ implemented in RELION^35^. CTF estimation was done using Gctf^36^ on non-dose-weighted micrographs. Template-free auto picking with the Laplacian-of-Gaussian (LoG) filter in RELION yielded ∼1.5 million particles, which were extracted in bin4. A 3D reconstruction was obtained after a few rounds of 2D Classification using a previously solved in-house apo structure of NavPas as the initial model. The resulting structure was then used for two rounds of 3D classification and 3D refinement on all the picked particles. Particles were then un-binned and subject to further rounds of 3D classification, resulting in 134 K particles. In parallel to this, template-based auto picking was also performed to give ∼2.9 million particles. Like LoG picked particles these were extracted in bin4, followed by two rounds of 2D classification and one round of 3D classification followed by un-binning. At this stage ∼110K particles were un-binned and further processed by 3D classification ultimately yielding ∼64K particles. The two particle stacks i.e., template based, and LoG picked were then joined and duplicate particles within 30 Å were removed. Such an approach of picking using two different methods and merging followed by duplicate removal has been shown to produce a larger number of particles that can ultimately lead to a better reconstruction^37^. This final stack contained ∼182K particles. After 3D refinement the particle stack was exported from RELION to Cryosparc and an additional round of 3D classification was followed by non-uniform refinement to produce a final map at 3.9 Å resolution generated with ∼164K particles as judged by the gold-standard Fourier shell correlation (GSFSC) = 0.143 criterion in CryoSparc^38^.

### Model building

The cryoEM structure of Apo NavPas (PDB code: 5X0M) and the NMR structure of LqhαIT (PDB codes: 1LQH) were fitted into the EM map using UCSF Chimera^39^. The model was refined in PHENIX^40^ using secondary structure and geometry restraints to prevent over-fitting. Model building was performed in COOT^41^.

### MD Simulation

The cryo-EM structure of the α-scorpion toxin-bound cockroach Nav channel (6nt4)^20^ was downloaded from the Orientations of Proteins in Membranes (OPM) database^42^. The in-house NavPas structure was aligned to the pdb: 6NT4 and processed with CCG MOE protein preparation, assigning standard amino acid protonation states at pH 7.4 and optimization of the hydrogen bonding network^43^ A POPC lipid bilayer was then built around the NavPas channel using the PACKMOL-Memgen utility,^44^ specifying a minimum distance of 10 Å between the receptor and box edge and a water layer thickness of 15 Å with NaCl salt concentration of 150 mM. This resulted in 686 lipid molecules, 77 473 water molecules, 208 sodium cations, and chloride anions. A single sodium cation was also inserted into the ion channel pore.

The membrane-embedded NavPas channel was first equilibrated for 100 ns using Amber20 and PMEMD CUDA^45–48^. The channel and toxin were modeled with ff19SB^49^, with water represented by TIP3P^50^, POPC lipids with Lipid21^51–54^, counter ions with JC parameters^55^ and sugars with GLYCAM06^56^ respectively. The system underwent initial minimization for 1000 steps using sander, employing the steepest descent method for the first 500 steps and the conjugate gradient method^57^ for the remaining steps. Subsequently, a longer minimization step of 10000 steps was conducted using PMEMD CUDA, with the first 5000 steps utilizing the steepest descent method and the remaining steps used the conjugate gradient method.

The system was then gradually heated from 0 to 100 K over a 5 ps constant volume run, employing Langevin dynamics^58^, with 10 kcal/mol/Å^2^ restraints on all channel, toxin, and lipid atoms. Afterward, the volume was allowed to change freely, and the temperature was raised to 298 K with a Langevin dynamics with anisotropic Berendsen control (γ = 1 ps^−1^) applied for 100 ps to maintain pressure around 1 atm 57 applied by coupling the periodic box with a time constant of 2 ps. The system was then run in the NPT ensemble with semi-isotropic Berendsen pressure coupling with the X- and Y-dimensions coupled, and the Z-dimension allowed to change freely for 1 ns with restraints on 5 kcal/mol/Å^2^ applied to channel and toxin backbone atoms. Another 1 ns of NPT simulation followed, with restraints reduced to 5 kcal/mol/Å^2^ on channel and toxin carbon-alpha atoms. Finally, all restraints were removed, and the system underwent a 100 ns simulation in the NPT ensemble. The system was then moved into a production simulation of 200 ns NPT with identical settings, except for the use of the Monte Carlo barostat for pressure regulation^59^. This entire equilibration and production protocol were iterated ten times with different random starting velocities.

All RMSD, RMSF, distance and angle analyses were performed on the resulting trajectories using CPPTRAJ^60^. Interaction counting was performed using the getcontacts utility^61^. All simulation results are presented as an average with standard deviation across the ten independent repeats.

## Acknowledgments

This work was supported in part by the Novartis Discovery Postdoctoral Fellowships (to C.J.S. and S.P.). The authors acknowledge Chad Vickers (Novartis Biomedical Research) for helpful scientific discussion and editorial input. Fig. 2A was produced using the UCSF Chimera package from the Resource for Biocomputing, Visualization, and Informatics at the University of California, San Francisco (supported by NIH P41 RR-01081).

## Author Contributions

S.P. and W.A.W. devised the project and designed the experiments. CryoEM data was collected by M.K. and processed by S.P. LqhαIT was purified by C.J.S., MD simulations were performed and analyzed by C.J.D. The manuscript was written by S.P., W.A.W., J.W., and C.J.S. with inputs from all authors.

## Data availability

The atomic coordinates and EM map for the NavPas-LqhαIT complex have been deposited in the Protein Data Bank (www.rcsb.org) and EMDB (www.ebi.ac.uk/pdbe/emdb/) with accession codes 8VQC and EMD-43438, respectively.

## Competing interests

All authors are/were employees of Novartis.

## Figure Legends

**Supplementary Fig 1. Purification and CryoEM of the LqhαIT-NavPas complex**. (A) Size exclusion chromatography trace of toxin-channel complex. (B) Fractions from the trace between the two dashed lines were run on SDS-PAGE gel and visualized by Coomassie staining, the peak fraction indicated by a red asterisk was concentrated and used for grid preparation. (C) GSFSC curves for the final reconstruction using various mask parameters. (D) Final reconstruction colored by local resolution. (E) A plot of pore radii along the Sodium ion permeation path as calculated by HOLE^65^, for the apo and LqhαIT-bound channel.

**Supplementary Fig 2. Comparison of Aah2-VSDIV-NavPas and LqhαIT-NavPas complex structures**. (A) Superimposition of Aah2-VSD-IV-NavPas (PDB: 6nt4)) and LqhαIT-NavPas complex structures with LqhαIT in orange, Aah2-Nav1.7-VSD-IV-NavPas chimera complex in light blue, VSD-IV and CTD in LqhαIT-bound NavPas in cyan and chartreuse-green, respectively, rest of NavPas is colored grey. (B) Superimposition of channel bound toxin structures showing broad similarities within α-helix and β-sheets with differences occurring in the loop and C-terminal regions. Coloring scheme same as A. (C) **Aah2-VSDIV-NavPas** interactions. Aah2 is colored light blue, Nav1.7-VSD-IV is colored lime green while the rest of the channel is colored grey. (D) Sequence alignment of LqhαIT and Aah2. Residues interfacing the protein or glycan part of the channel (with interface areas >25 Å^2^ are marked by green or blue asterisks, respectively. (E) Lqh2 mutagenesis data plotted alongside buried surface area (BSA) for Aah2. Lqh2 here acts as a surrogate for Aah2 with (only two different amino acids corresponding to 96% sequence identity) for which mutagenesis data is available from Kahn et al^27^. BSA was calculated for the Aah2-Nav1.7-VSD-IV-NavPas complex (PDB: 6nt4) using PISA^64^. The X axis indicates amino acid mutation and the Y axis on the right shows the corresponding change in EC50 ratio of mutant over wildtype Lqh2 plotted as orange dots. The Y axis on the left indicates buried surface area as observed for these residues in the Aah2-Nav1.7-VSD-IV-NavPas complex structure. Bars represent area buried by the glycan moiety (blue) or channel residues (dark green).

**Supplementary Fig 3**. (A) Sequence conservation for the DI pore glycan site at Asn330 (NavPas numbering). (B) Comparison of Nav channel-scorpion α toxin binding modes. Cartoon representations of VSD-IVs and toxins, while glycans are shown as sticks. Lqh3 interacts with a more peripheral epitope on Nav1.5 VSD-IV compared to Aah2 and LqhαIT, which bind in close proximity to corresponding glycans atop their DI-PDs. Distances between the 2^nd^ NAG residues and the center of mass of respective toxins are shown.

**Supplementary Fig 4**. (A) Changes at the S4-S5 linker of NavPas upon LqhαIT binding. Apo NavPas colored yellow and NavPas bound to LahαIT colored cyan. R5 residues at the base of S4 are shown as sticks for reference. A black line indicates displacement of the residue Arg 1283 at the beginning of S5 by 3.4 Å.

**Movie 1**. Activation of VSD-IV within the NavPas channel is depicted through a trajectory. NavPas bound by LqhαIT structure (in cyan) and the apo state structure (in yellow). The R5 residues situated at the base of S4 are represented as sticks for reference, while S3 is omitted for clarity.

