## Supplementary figures and images for "Scorpion α-toxin LqhαIT specifically interacts with a glycan at the pore domain of voltage-gated sodium channels"

### Supplementary fig. 1

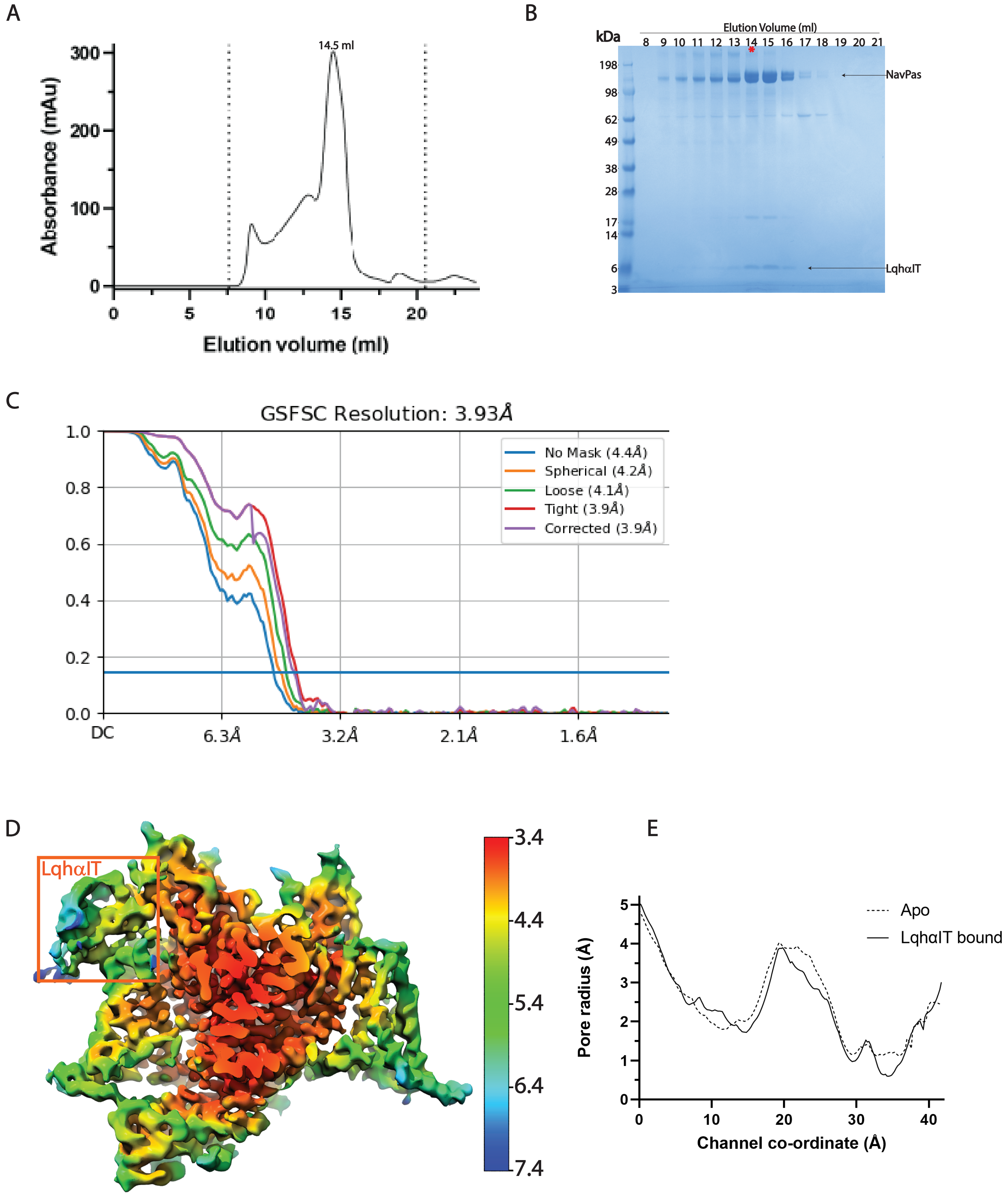


Supplementary Fig 1

### Supplementary fig. 4

A

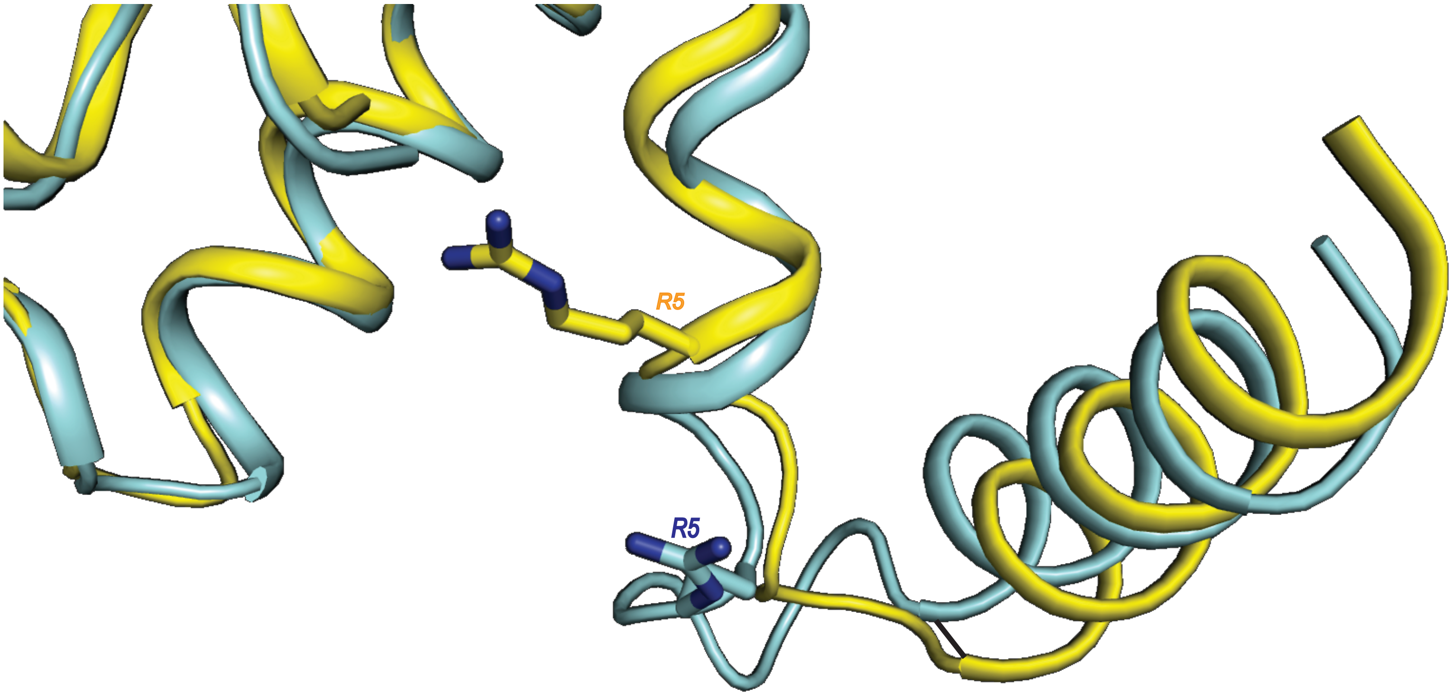


Supplementary Fig 4
