## Supplementary fig. 2 for "Scorpion α-toxin LqhαIT specifically interacts with a glycan at the pore domain of voltage-gated sodium channels"

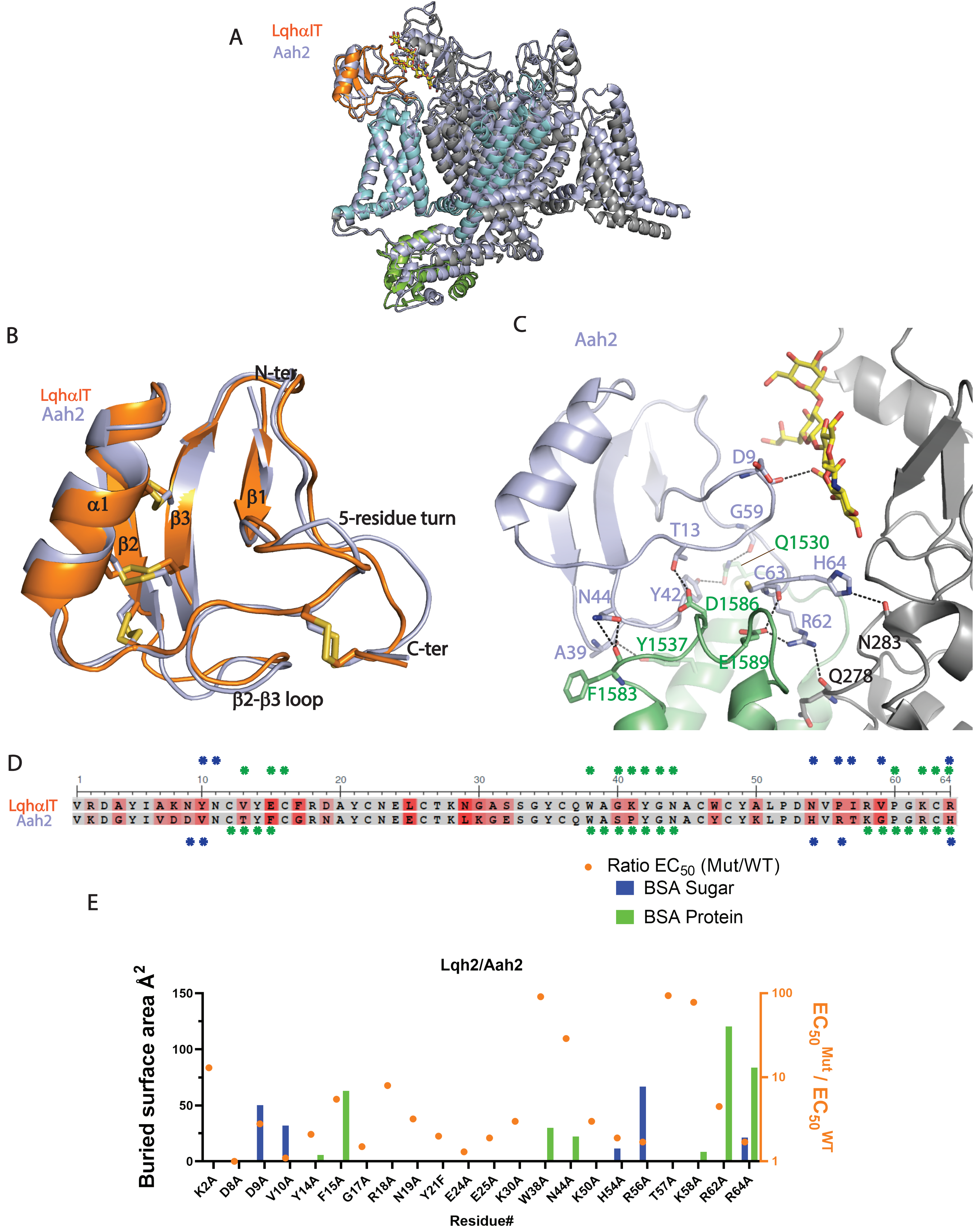


Mutational data for Lqh2, BSA Protein/Sugar derieved from Aah2-Nav1.7-NavPaS chimeric complex structure.Lqh2 is almost identical to Aah2 with ~96% identity

Supplementary Fig 2
