## Supplementary fig. 3 for "Scorpion α-toxin LqhαIT specifically interacts with a glycan at the pore domain of voltage-gated sodium channels"

A

| | 330 |
|---|---|
| **NavPas** | YPLCG**N**SSGAG |
| **NavBg** | VPLCG**N**SSGAG |
| **Nav1.1** | ALLCG**N**SSDAG |
| **Nav1.2** | ALLCG**N**SSDAG |
| **Nav1.3** | PLLCG**N**GSDAG |
| **Nav1.4** | ALLCG**N**SSDAG |
| **Nav1.5** | VLLCG**N**SSDAG |
| **Nav1.6** | PLLCG**N**SSDAG |
| **Nav1.7** | ALLCG**F**STDSG |
| **Nav1.8** | PLLCG**N**DSGSD |
| **Nav1.9** | FKMCG**I**WMGNS |

B

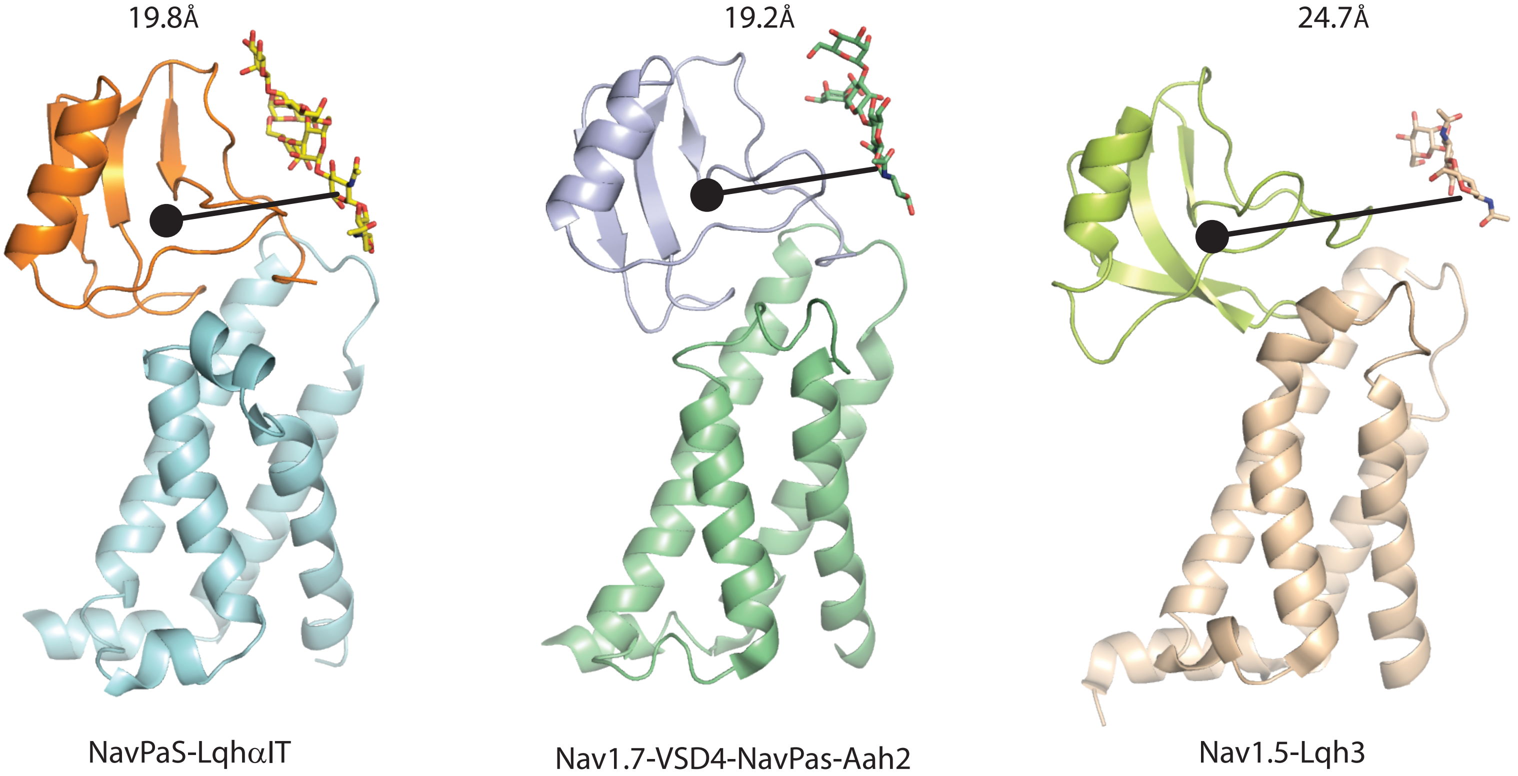


Supplementary Fig 3
